# COP9 signalosome is an essential and druggable parasite target that regulates protein degradation

**DOI:** 10.1101/2020.03.24.004531

**Authors:** Swagata Ghosh, Laura Farr, Aditya Singh, Laura-Ann Leaton, Jay Padalia, David Sullivan, Shannon Moonah

## Abstract

Understanding how the protozoan protein degradation pathway is regulated could uncover new parasite biology for drug discovery. We found the COP9 signalosome (CSN) conserved in multiple pathogens such as *Leishmania, Trypanosoma, Toxoplasma*, and used the severe diarrhea-causing *Entamoeba histolytica* to study its function in medically significant protozoa. We show that CSN is an essential upstream regulator of parasite protein degradation. Genetic disruption of *E. histolytica* CSN by two distinct approaches inhibited cell proliferation and viability. Both CSN5 knockdown and dominant negative mutation trapped cullin in a neddylated state, disrupting UPS activity and protein degradation. In addition, zinc ditiocarb (ZnDTC), a main metabolite of the inexpensive FDA-approved alcohol-abuse drug disulfiram, was active against parasites acting in a COP9-dependent manner. ZnDTC, given as disulfiram-zinc, had oral efficacy in clearing parasites in vivo. Our findings provide insights into the regulation of parasite protein degradation, and supports the significant therapeutic potential of COP9 inhibition.

**Summary sentence:** Parasite-encoded COP9 signalosome is an essential upstream regulator of ubiquitin-proteasome mediated protein degradation, and shows significant potential as a therapeutic target.

## INTRODUCTION

Protein turnover, which is the balance between protein synthesis and protein degradation, is essential for life. The Ubiquitin-Proteasomal System (UPS) is conserved in all eukaryotes including protozoans, and is responsible for the vast majority of protein degradation within the cell. Cullin-based ubiquitin ligases catalyze the ubiquitination of proteins destined for proteasomal degradation. Disruption of proteasomal activity results in accumulation of unwanted and toxic proteins, ultimately leading to cell death (*1*).

Protozoan parasites, such as *Trypanosoma, Entamoeba, Leishmania, and Toxoplasma*, present a major threat to global public health, and contribute significantly to morbidity and mortality worldwide. Antibiotic treatment is essential for managing patients infected with these parasites. That said, new therapies are urgently needed given the lack of effective vaccines, drug resistance, limited efficacy and toxicity associated with current treatment (*2-4*). Given that the UPS pathway is essential for cell survival, inhibition of the proteasome has emerged as an attractive anti-parasitic target (*5-9*). However, the regulation of UPS-mediated protein degradation in clinically important protozoans remains poorly understood. Understanding the regulatory pathways involved in proteasomal degradation could, therefore, lead to new therapeutic opportunities for patients with these difficult-to-treat infections, either as monotherapy or in combination with established parasite proteasomal inhibitors.

Here, we investigated COP9 (Constitutive photomorphogenesis 9) signalosome function in disease-relevant protozoa using *Entamoeba histolytica* as the model parasite. *E. histolytica* is a protozoan parasite that is a leading cause of severe diarrhea in the world’s poorest communities. There is no vaccine and only one class of drugs (nitroimidazoles) available to effectively treat invasive forms of disease. This is a major concern as we are ill-prepared and left with no option if resistance or intolerable side effects develops (*4, 10-12*). COP9 signalosome was found to be encoded by other protozoans such as *Leishmania, Trypanosoma*, and *Toxoplasma*, in addition to *E. histolytica*. We uncover *E. histolytica* COP9 signalosome as an essential upstream regulator of the parasite UPS protein degradation pathway. The zinc-ditiocarb complex, a major metabolite of disulfiram, inhibited the *E. histolytica* COP9 activity, highlighting the potential for repurposing disulfiram as an anti-parasite agent.

## RESULTS

### Characterization of parasite-produced COP9 signalosome

The COP9 signalosome (CSN), is a multi-subunit protein complex that is conserved across animals and plants, but remains largely uncharacterized in pathogenic protozoan parasites (*13-16*). We identified subunit 5 of COP9 signalosome (CSN5) in *E. histolytica*, which was found to be highly conserved in other pathogenic protozoans such as *Leishmania, Trypanosoma, Toxoplasma, Acanthamoeba and Naegleria*. The parasite CSN5 contained the JAMM (JAB1/MPN/Mov34 metalloenzyme) motif, which is the zinc metalloprotease site consisting of the amino acid sequence HXHX_7_SX_2_D (Fig. 1A).

**Fig. 1.**
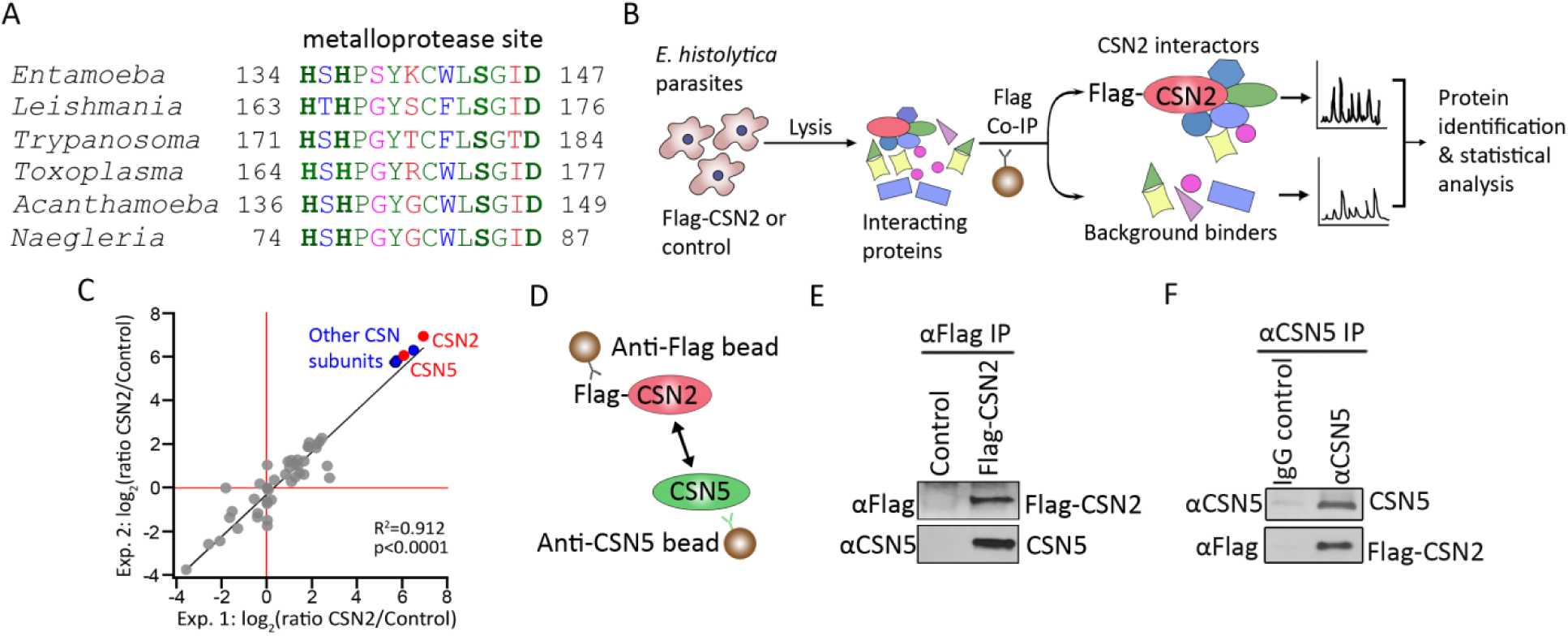
Characterization of parasite-encoded COP9 signalosome. (**A**) Multiple sequence alignment of the COP9 signalosome subunit 5 (CSN5) metalloprotease site from medically important protozoan parasites. Identical (green), conserved (blue), semi-conserved (pink) and non-conserved residues (red). (**B**) Experimental approach for identifying CSN2 interacting proteins from *E. histolytica* cells by co-immunoprecipitation (Co-IP) followed by mass spectrometry. (**C**) Scatter plot and Pearson’s correlation analysis of log2 ratios label-free quantification intensities for proteins identified by mass spectrometry in anti-Flag co-immunoprecipitations from *E. histolytica* cells expressing Flag-CSN2 compared to the empty vector control. CSN2 and CSN5 subunits (red), putative COP9 subunits (blue), and non-COP9 subunits (grey). (**D**) Diagram of experimental procedure for validating CSN5 and CSN2 interaction. (**E** and **F**) Reciprocal co-immunoprecipitation with anti-Flag (E) and anti-*E. histolytica* CSN5 (F) antibodies show interaction between CSN5 and CSN2.

Next we investigated whether the CSN5 protein was a true component of the parasite COP9 signalosome complex by determining the interaction between CSN5 and other subunits. Among the subunits that make up the COP9 signalosome complex, CSN2 and CSN5 are the most conserved subunits (*15, 17*). Therefore, we investigated the interaction between the parasite CSN2 and CSN5. We identified the CSN2 gene and created a Flag tagged CSN2 expressing ameba cell line, followed by affinity pulldown with anti-Flag antibody (fig. S1 and Fig. 1B). The immunoprecipitated endogenous proteins were first analyzed by quantitative mass-spectrometry, which revealed CSN2 and CSN5 subunits, along with other CSN subunits as the most enriched co-purified proteins (Fig. 1, B and C). In addition, protein–protein interaction screening using immunoaffinity purification of amebic cell lysate with specific anti-CSN5 antibody followed by mass spectrometry analysis revealed similar results (fig. S2). Although we did not co-purify CSN4 and 8, we identified the putative genes in the *E. histolytica* genome. CSN5 and CNS2 protein-protein interaction was confirmed by reciprocal co-immunoprecipitation (Fig. 1, D to F). These findings suggest that the CSN5 identified in the *E. histolytica* is a bona fide component of the parasite COP9 signalosome complex.

### COP9 signalosome is necessary for parasite protein degradation

To characterize the role of COP9 signalosome in *E. histolytica* biology, we first used a genetic knockdown approach. Because subunit 5 forms the catalytic center of the COP9 signalosome (CSN5) (*14*), we evaluated the function of the parasite COP9 by CSN5 gene silencing using an established high-efficiency inducible RNA interference technology (*18, 19*). CSN5 knockdown was validated by immunoblotting with anti-*E. histolytica* CSN5 antibody, and a marked decrease in the CSN5 proteins level was observed in the knockdown cells compared to the controls (fig. S3A). Next we performed a cell viability assay and found that CSN5 knockdown resulted in a proliferation defect and eventual cell death within 36 hours when compared with the empty vector control (Fig. 2A).

**Fig. 2.**
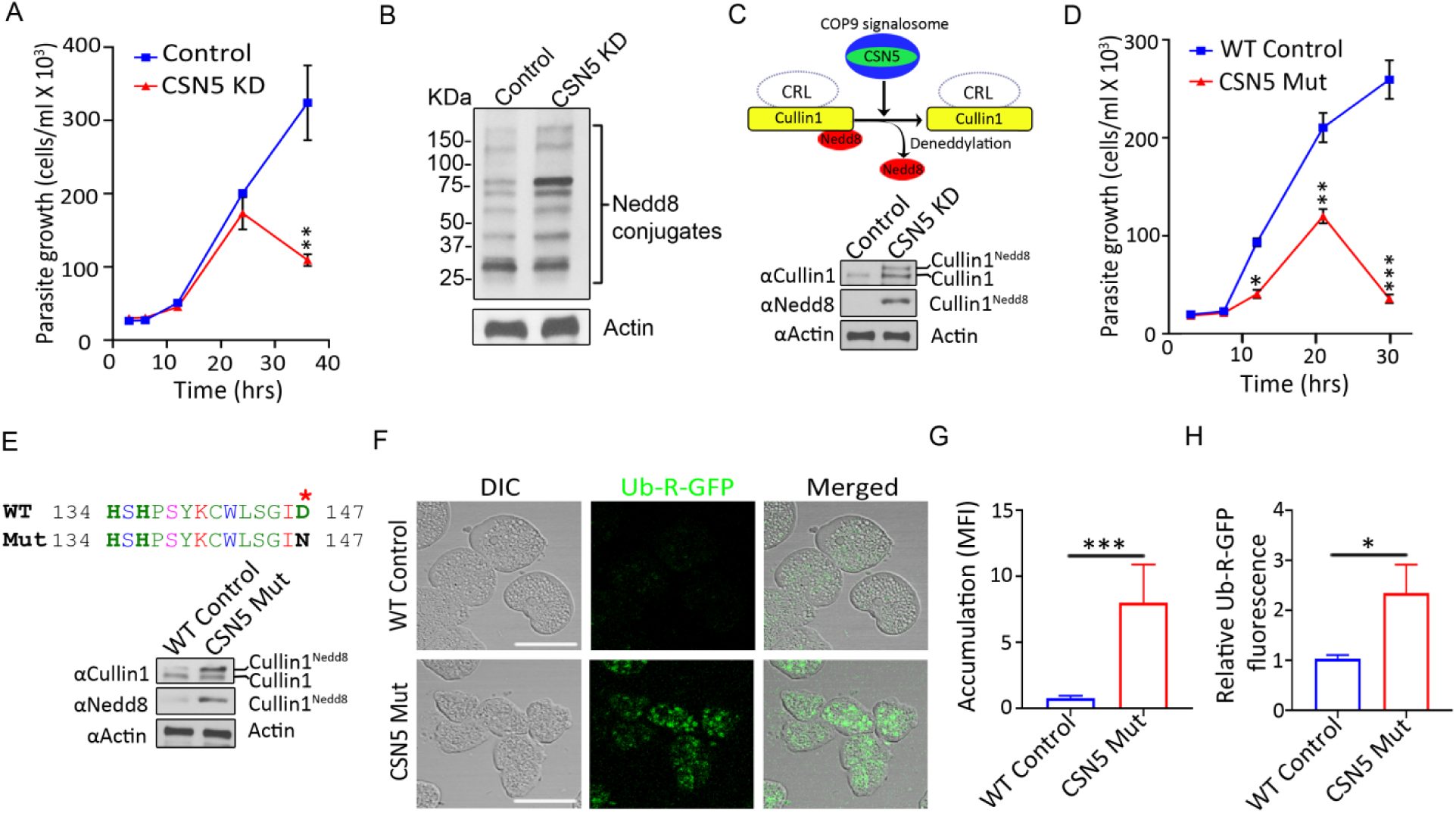
COP9 signalosome activity is essential for *E. histolytica* protein degradation. (**A**) Cell proliferation assay showing the effect of CSN5 knockdown on *E. histolytica* viability. (**B**) Immunoblot with anti-*E. histolytica* Nedd8 antibody demonstrating accumulation of neddylated proteins in CSN5 knockdown cells compared to the empty vector control. Actin was used as a loading control. (**C**) Schematic of cullin1 deneddylation by the COP9 signalosome. Accumulation of neddylated cullin1 in CSN5 knockdown cells compared to empty vector control (24 h). Immunoblot of amebic lysate with anti-*E histolyica* cullin1 antibody. Immunoprecipitated cullin1 was analyzed by immunoblotting using anti-*E. histolytica* Nedd8 antibody. (**D**) Dominant negative effect of the catalytically inactive CSN5 mutant. Cell proliferation assay evaluating the effect of a single amino acid mutation (D147N) within the CSN5 metalloprotease site on parasite viability. (**E**) Immunoblot analysis with anti-*E. histolytica* cullin1 and Nedd8 antibodies on CSN5 WT and mutant (D147N) overexpressing cells showing levels of neddylated cullin1. (**F** and **G**) CSN5 dominant negative mutant expression impairs protein degradation. Confocal images and quantification of WT and CSN5 mutant expressing live cells accumulating the protein degradation substrate ubiquitin-arginine-GFP (Ub-R-GFP). Mean fluorescence intensity, MFI. (**H**) Fluorometric assay of Ub-R-GFP accumulation. CSN5 mutant results in GFP accumulation. Data represent mean ± SD of quintuples from one experiment and are representative of three independent experiments. \**P* < .05, ** *P* < 0.01, *** *P* < 0.001, two-tailed *t* test.

Deneddylation involves the removal of the ubiquitin-like protein Nedd8 from Nedd8-conjugated (neddylated) proteins (*14, 20*). Given that COP9 complex has deneddylation activity in non-parasitic eukaryotes and Nedd8 proteins are conserved in parasites (*21, 22*), we hypothesized that the parasite COP9 has deneddylating function. First, we examined the effect of CSN5 depletion on global protein neddylation and found that knocking down CSN5 resulted in an increase in neddylated proteins compared to the controls (Fig. 2B), which suggested that the COP9 deneddylation activity is inhibited upon CSN5 knockdown.

Cullins are the best characterized neddylated COP9 substrates in other eukaryotes (*23*). A conserved cullin lysine residue forms an isopeptide bond with the carboxy-terminal Gly-76 of the Nedd8 protein (*24*). We have previously identified an *E. histolytica* cullin1 protein (*16*) which has the conserved neddylation site (fig. S4). Immunoblot analysis revealed the most intense neddylated band after CSN5 disruption being above the 75 KDa mark, which includes neddylated cullin1 (Fig. 2B, fig. S3E), which can sometimes run lower than its predicted molecular weight (*25*). Therefore, we tested if cullin1 neddylation is altered upon CSN5 knockdown, and found that knockdown cells accumulated neddylated cullin1 by immunoblot analysis using antibodies specific for *E. histolytica* cullin1 and Nedd8 (Fig. 2C). This is consistent with a recent cancer biology study that showed a specific COP9 inhibitor caused accumulation of neddylated cullin prior to cancer cell proliferation defects (*26*). Thus, disrupting *E. histolytica* COP9 by CSN5 knockdown resulted in accumulation of cullin1 in its neddylated state.

In addition to the knockdown approach, we utilized dominant negative mutant expression as an independent strategy to inhibit endogenous protein function. Overexpression of enzymatically dead mutant proteins result in dominant negative effect. This method has been successfully used to explore gene function in *E. histolytica* biology (*27*). A single amino acid substitution, Asp 147→Asn 147 (D147N), was generated in the metalloprotease site of CSN5, which abolishes its zinc binding ability rendering it catalytically inactive (fig. S3B and S5A). Consistent with CSN5 knockdown, cells expressing the mutant protein had severe cell proliferation and viability defects (Fig. 2D).

Cullin based RING E3 ubiquitin ligases (CRLs) tag cellular proteins with ubiquitin (Ub) chains for their subsequent degradation by the proteasome, and CRL activity is dependent on deneddylation of cullins (*13, 15, 28*). Similar to the CSN5 knockdown, CSN5 mutant expression resulted in accumulation of cullin1 in its neddylated form (Fig. 2E, fig. S3C). Next, we investigated the effect of parasite COP9 disruption on UPS-dependent protein turnover using an *E. histolytica* strain that constitutively express a GFP reporter substrate of UPS-mediated protein degradation, ubiquitin-arginine-GFP (Ub-R-GFP). We found that mutant CSN5 overexpression disrupted protein degradation resulting in accumulation of GFP as measured by quantitative confocal imaging (Fig. 2, F and G) and fluorometric assay (Fig. 2H). Taken together, our data indicate that the parasite COP9 signalosome is essential for protein degradation.

### ZnDTC is active against *E. histolytica* in a COP9-dependent manner and phenocopy CSN5 genetic disruption

Our finding that COP9 signalosome function was essential for *E. histolytica* cell survival led us to test its druggability. Since COP9 essential deneddylation activity is embedded within the metalloprotease site of the CSN5 subunit, we screened known metalloprotease inhibitors and chelating agents for their ability to dock onto CSN5 using virtual screening by molecular docking (*29*). Among the hits of potential inhibitors was zinc-ditiocarb (ZnDTC) (fig. S5A). ZnDTC was of particular interest because it is a metabolite of the drug disulfiram (Fig. 3A). Disulfiram has been approved by the FDA since 1951 to treat alcohol dependence with safety and pharmacokinetics profile already established (*30*).

**Fig. 3.**
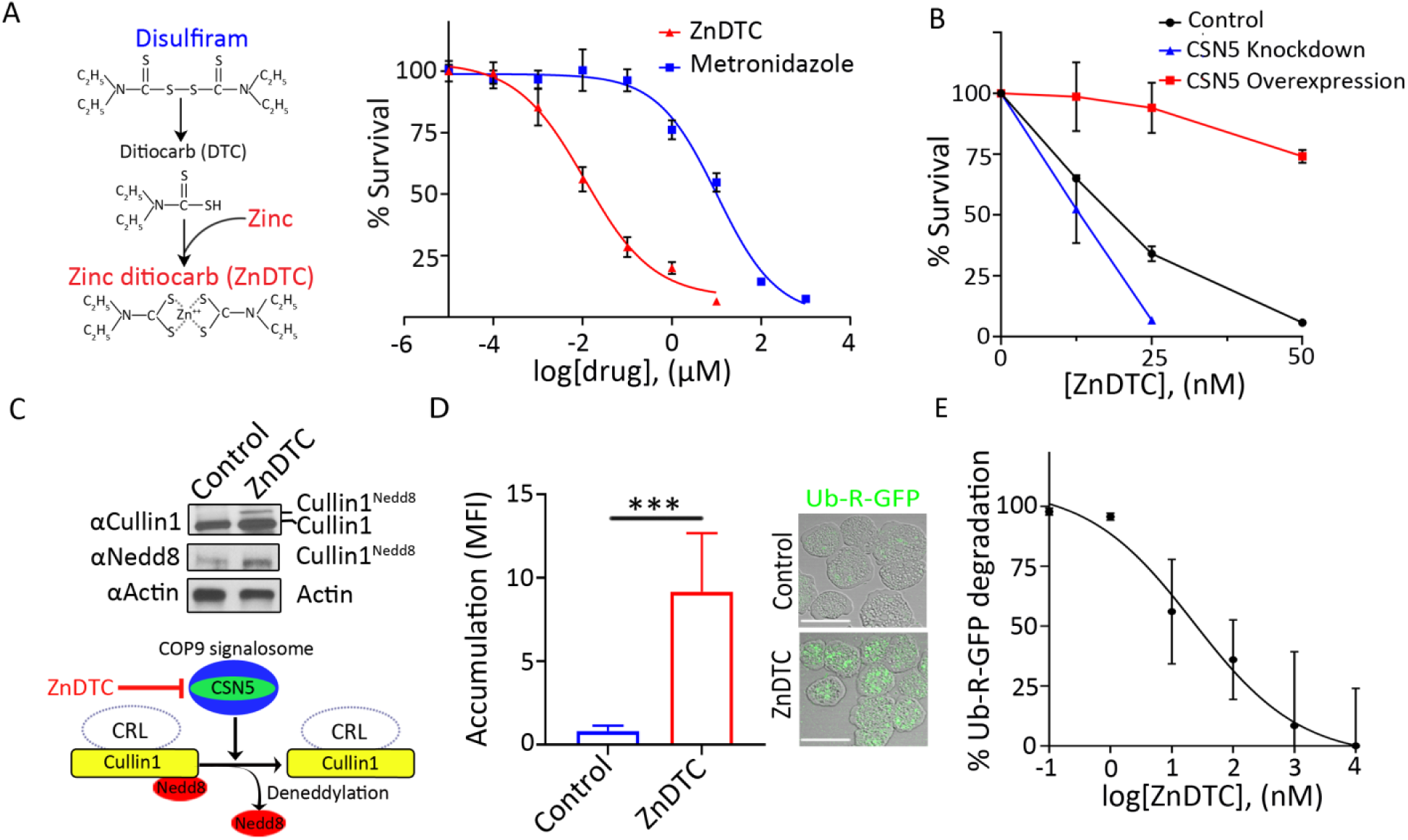
Pharmacological inhibition of COP9 by ZnDTC phenocopy CSN5 genetic disruption. **(A**) Illustration of zinc-ditiocarb (ZnDTC) formation from disulfiram and zinc. Dose response curve showing amebicidal effect of ZnDTC and metronidazole after 48 hours of incubation at indicated doses. EC_50_ for ZnDTC (11.21 ± 6.39 nM) and metronidazole (10.26 ± 6.06 µM) was calculated from nonlinear robust regression fit of the dose response curves. (**B**) Target validation using viability assay of empty vector control, CSN5 knockdown and CSN5 overexpression parasites incubated with increasing doses of ZnDTC. CSN5 overexpression reduces sensitivity to ZnDTC while CSN5 knockdown increases sensitivity to ZnDTC. (**C**) Immunoblot analysis for neddylated cullin1 with anti-*E. histolytica* cullin1 and Nedd8 antibodies on ZnDTC treated cells. (**D**) Quantification of live cells accumulating the protein degradation substrate, Ub-R-GFP, in ZnDTC treated cells with representative confocal micrographs. (**E**) Fluorometric assay of Ub-R-GFP degradation. ZnDTC inhibits Ub-R-GFP degradation in a dose dependent manner. Data represent mean ± SD of quintuples from one experiment and are representative of three independent experiments. \*\*\**P* < .001, two-tailed *t* test.

To evaluate whether ZnDTC had anti-amebic activity, we determined the dose response curve for the effect of ZnDTC on parasite viability. The compound was effective against parasites at nanomolar concentration with an EC_50_ of 11.21 ± 6.39 nM (Fig. 3A), which is below the serum and tissue level achieved in individuals on disulfiram therapy at recommended doses (*31, 32*). The EC_50_ of metronidazole, the current drug of choice to treat amebiasis, was found to be 10.26 ± 6.06 µM (Fig. 3A). This is in keeping with previous reports for metronidazole EC_50_ ranging from 5 to 20 µM (*11, 12*).

Drug resistance by gene overexpression is a useful approach for target identification and validation (*33, 34*).Therefore, to test if CSN5 is the target of ZnDTC we examined the susceptibility of cells overexpressing CSN5. Overexpression of CSN5 resulted in increased resistance to ZnDTC treatment (fig. S3B, Fig. 3B, fig. S5B), providing evidence that CSN5 is targeted by ZnDTC in *E. histolytica* parasite. As expected, concentrations of 100 nM and greater resulted in significant growth defect (fig. S5B), which occurs when drug concentrations exceed overexpression capacity (*33*). In support of these findings, CSN5 knockdown rendered cells more sensitive to ZnDTC (fig. S3A, Fig. 3B).

To further evaluate whether ZnDTC acts in a COP9 dependent manner, we examined cullin1 deneddylation and UPS activity in drug-treated cells compared to controls. We found that cells treated with ZnDTC phenocopy genetic disruption of CSN5 by trapping cullin1 in a neddylated state (Fig. 3C, fig. S3D, fig. S3F), and inhibited protein degradation (Fig 3, D and E). Collectively, these findings imply that ZnDTC exerts anti-amebic activity by disrupting *E. histolytica* COP9 signalosome-regulated proteolysis.

### ZnDTC has potent activity against *E. histolytica* in a mouse model that mirrors human infection

We next tested the therapeutic effects of zinc ditiocarb in vivo. In the body, disulfiram is rapidly metabolized to ditiocarb (DTC) which in the presence of metal ions such as zinc, forms zinc-ditiocarb complex (ZnDTC). DTC-metal complexes have a relatively long half-life and are widely distributed throughout the body, including the gastrointestinal tract. Therefore, in order to achieve adequate levels, ZnDTC is safely given in vivo as oral disulfiram plus zinc gluconate (*31, 32, 35-38*). *Entamoeba histolytica* causes an inflammatory diarrhea termed amebic colitis (*10*). We used a mouse model that simulates human amebic colitis for in vivo studies. Mice were infected with luciferase-expressing parasites and infection was monitored by live bioluminescence imaging. Infected mice received a 5 day treatment course based on the minimum recommended treatment duration for human amebiasis (*10*). Similar to the anti-amebic drug metronidazole (*39, 40*), we started to observe parasite clearance after 2 days of ZnDTC therapy (Fig. 4A). Consistent with our live imaging findings, all ZnDTC-treated mice were culture negative at the end of the 5-day treatment course, compared to none in the untreated group (Fig. 4, A and B, fig. S5C). Histopathological examination and immunohistochemical staining using specific anti–*E. histolytica* antibodies revealed numerous parasites in untreated mice, absent in the treated mice (Fig. 4C). In addition, ZnDTC significantly reduced the destructive inflammatory response, and tissue damage as measured by tissue myeloperoxidase (MPO) levels and histological score (Fig. 4, C to E). These findings support an in vivo antiparasitic effect.

**Fig. 4.**
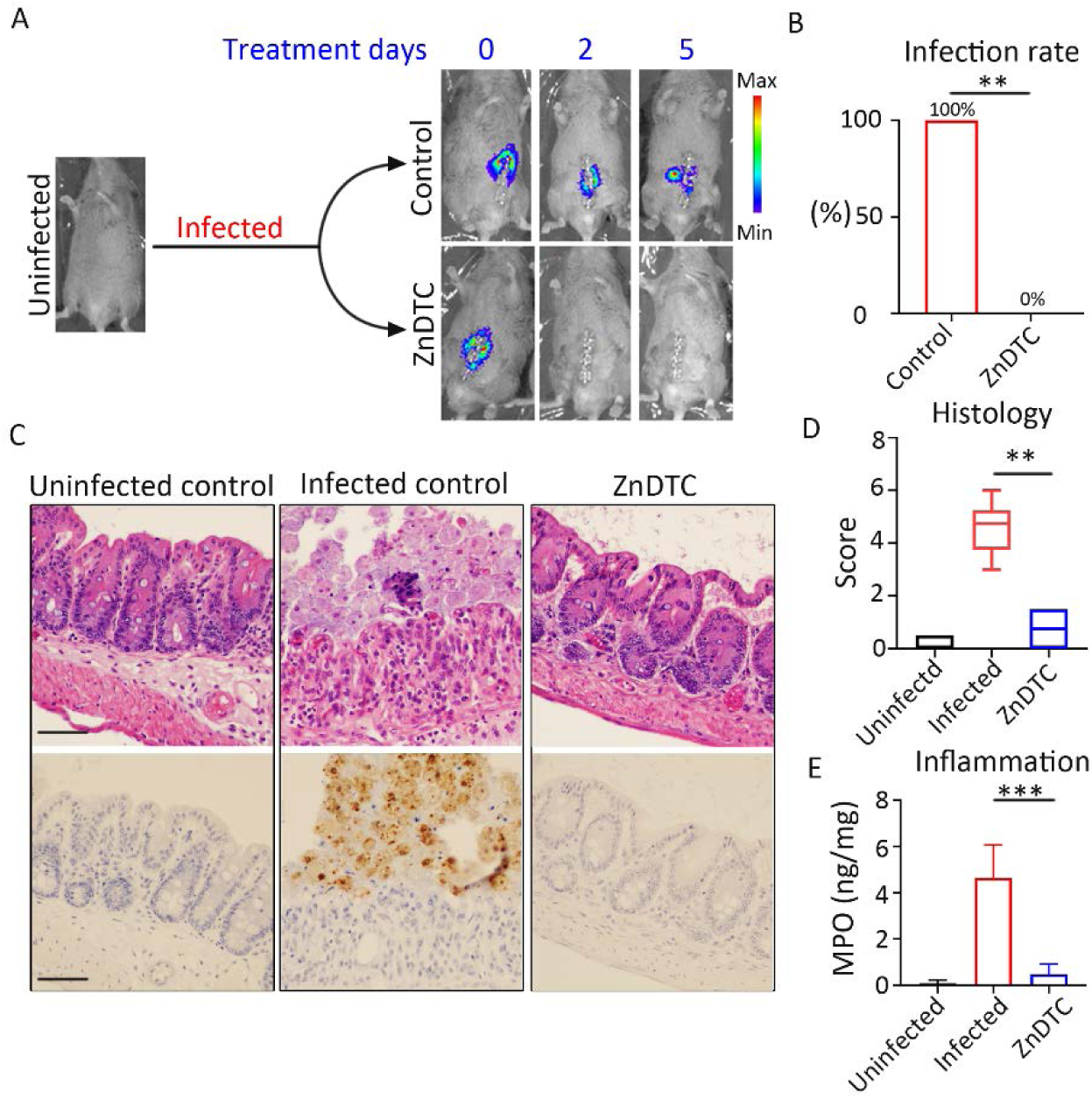
ZnDTC has potent activity against *E. histolytica* in a preclinical animal model of amebic colitis. (**A**) Representative live bioluminescent images of mice infected with luciferase-expressing *E. histolytica* during the treatment period. (**B**) Infection rate measured by ameba culture of cecal content. (**C**) Representative H&E staining and immunohistochemical analysis of the cecum of infected mice after 5 days of treatment. Specific anti–*E. histolytica* macrophage migration inhibitory factor antibody was used for immunohistochemical staining of trophozoites (brown). Numerous parasites in the infected control, absent in the ZnDTC treated mice. Scale bar, 50μm. (**D**) Histology score. (**E**) Reduced levels of the MPO marker of intestinal inflammation in mice treated with ZnDTC. Data represent mean ± SD (*n* = 6 mice per group). \*\**P* < .01, \*\*\**P* < .001, Fisher’s exact test and Mann-Whitney *U* test.

## DISCUSSION

Protozoan parasites continue to pose serious threats to health and contribute significantly to morbidity and mortality worldwide. Those living in poverty-stricken settings are disproportionately affected. Limited treatment options, poor efficacy, drug resistance, toxicity and expense contribute to the poor health outcomes associated with current therapies. Parasitic protozoan diseases are in dire need of new drugs. The ubiquitin proteasome pathway is the main engine for parasite protein degradation and therefore regulates multiple essential biological processes. Hence, the ubiquitin proteasome pathway has received tremendous attention recently in the parasitology field as an attractive drug target (*1*). Yet, our understating of how the ubiquitin proteasome pathway is regulated in these disease-relevant parasites remain limited. Here, *E. histolytica* was used as a prototype to study the role of COP9 in protein degradation in a medically significant protozoan. We found that the COP9 signalosome was essential for parasite biology, as disruption of the parasite COP9 led to dysregulation of the ubiquitin proteasome pathway which impaired protein degradation and led to cell death.

All pathogenic protozoans express cullins and Nedd8 proteins. While the majority encode COP9, it appears some do not, including Plasmodium parasites. Plasmodium encodes hydrolases that are capable of cleaving Nedd8, and while important, these enzymes appear to lack the ability to deneddylate cullin (*22*). It remains possible that these parasites produce proteins with CSN5-like characteristics and activity, and we should be encouraged that with continued research we might be able to identify these proteins.

Disulfiram, also known as Antabuse, is an FDA-approved drug that has been used to treat alcohol dependence for several decades, with well-established pharmacokinetic properties, safety and tolerance (*30*). Disulfiram has shown varying activity against parasites, with an even greater antiparasitic potency observed when complexed to zinc (ZnDTC) (*41*). However, these observations were based mainly on *in vitro* studies, and the underlying mechanisms that explain the enhanced potency remain poorly understood. In this study, we showed that the disulfiram metabolite, ZnDTC, has anti-parasite effects *in vitro* and *in vivo*, and identified the COP9 signalosome as a drug target. These findings are attractive for several reasons. Firstly, disulfiram penetrates a wide range of tissues including the blood-brain barrier, and so holds potential use in the treatment of infections of the central nervous system caused by COP9 producing protozoans. Disulfiram is well-tolerated and has excellent bioavailability, therefore the drug can be given orally for extended periods. For example, the combination of oral zinc gluconate and disulfiram was well-tolerated for 53 continuous months with negligible side effects in a patient with metastatic cancer (*35*). Furthermore, disulfiram is a globally available, economical and low-cost drug, which could make it an affordable option for patients in low income countries (*37*). Our findings provide mechanistic insight into the antiparasitic activity of disulfiram, and establish *in vivo* efficacy, suggesting that disulfiram may be suitably poised for further drug development under repurposed indications. Established homologs present in humans can be an issue in drug development. That said, the ZnDTC safety profile plus success targeting parasite proteasome (*1*) provide reasons to be optimistic. In addition, COP9 signalosome inhibitors are being optimized to treat cancer (*26*), hence one could envision repurposing these drugs for use against parasitic protozoal diseases. The possibility of developing a disulfiram reaction is not anticipated to deter treatment adherence under potential repurposed indication, as there are many other safe and well-established, FDA-approved medications that interact with alcohol, and hence also require abstinence while on treatment without affecting compliance. The commonly prescribed antibiotic metronidazole, for example, which coincidentally belongs to the only drug class available for the treatment of invasive amebic disease, has been reported to lead to the development of an unpleasant and even life-threatening disulfiram-like reaction when alcohol is concomitantly consumed (*42, 43*). Despite this, metronidazole has become the treatment of choice for several parasitic and anaerobic infections and is used worldwide with good adherence (*43*).

In conclusion, we provide genetic and chemical evidence for COP9 signalosome as an anti-protozoan drug target. These findings advance in our understanding of parasite protein turnover, and the results can be potentially leveraged to create new therapeutic opportunities to address the unmet medical needs for individuals suffering from these neglected diseases.

## MATERIALS AND METHODS

### Study approval

All animal procedures were approved by the University of Virginia Institutional Animal Care and Use Committee. All animal studies were performed in compliance with the federal regulations set forth in the Animal Welfare Act, the recommendations in the Guide for the Care and Use of Laboratory Animals of the National Institutes of Health, and the guidelines of the University of Virginia Institutional Animal Care and Use Committee.

### Molecular cloning and protein purification

*E. histolytica* CSN5, cullin1 and Nedd8 genes, codon optimized for expression in *E. coli* were amplified with primers carrying 5′BamHI and 3′XhoI sites and cloned within BamHI and XhoI sites of pGEX-6P1 vector to generate GST-fusion proteins. To express MBP-fusion versions of these proteins, each of the three genes were sub-cloned within the 5′BamHI and 3′XhoI sites of pET28-MBP-TEV vector, a gift from Zita Balklava & Thomas Wassmer (Addgene plasmid # 69929). *E. coli* BL21(DE3) cells were used for expression and purification of GST and GST fusion proteins including *E. histolytica* CSN5, cullin1 and Nedd8. Recombinant protein expression was done using previously described protocol (*16*). MBP and MBP fused *E. histolytica* CSN5, cullin1 and Nedd8 proteins were expressed by induction with 0.1 mM isopropyl β-D –thiogalactoside (IPTG) for 4 hours at RT. Purification of GST and MBP fusion proteins were done using glutathione-sepharose (GE-Healthcare) and amylose resin (New England Biotechnologies), respectively. Amicon® Ultra-15 Centrifugal Filter Units (Millipore Sigma) were used to concentrate and buffer exchange proteins into 1x PBS. For tetracycline dependent overexpression and knockdown of CSN5 in *E. histolytica* cells, the CSN5 gene was amplified from genomic DNA and was cloned in the shuttle vector pEhHYG-tetR-O-CAT (*18*) in place of the CAT gene using KpnI and BamHI in either the sense or the antisense orientation, respectively. For overexpression of the mutant CSN5 in a tetracycline dependent manner, D147N mutation was introduced within the CSN5 overexpression construct (pEhHYG-tetR-O-CSN5 sense clone) by inverse PCR with 5’ phosphorylated primers and Platinum™ SuperFi™ DNA Polymerase (Thermo Fisher Scientific # 12351010) followed by Dpn1 digestion and self-ligation. The *E. histolytica* CSN2 gene was cloned into pKK-FLAG-TEV vector, a gift from Andrzej Dziembowski (Addgene plasmid # 105768; http://n2t.net/addgene:105768; RRID:Addgene_105768), using BamhI and XhoI sites. The Flag-CSN2 was then sub-cloned into the NheI and SmaI sites of pKT3 vector for its constitutive expression in amoeba. The UPS reporter construct Ub-R-GFP was obtained as a gift from Nico Dantuma (Addgene plasmid # 11938; http://n2t.net/addgene:11938; RRID:Addgene_11938) and sub-cloned into the pKT3 plasmid using the NheI and SmaI sites. The luciferase plasmid pHTP.luc was used to generate the luciferase expressing ameba.

### Parasite culture and transfection

*Entamoeba histolytica* strain HM1:IMSS trophozoites were grown at 37°C in TYI-S-33 medium. Parasites were transfected using Attractene (Qiagen). The transfectants were selected with hygromycin B for pEhHYG-tetR-O-CAT based plasmids and G418/neomycin for pHTP.luc and pKT3 based plasmids. The initial selections were done at a concentration of 9µg/ml antibiotic and gradually increased up to 20µg/ml except for the pHTP.luc plasmid which was finally selected with 50µg/ml G418. The parasites transfected with pEhHYG-tetR-O-CAT empty or CSN5 constructs (overexpression/sense, knockdown/antisense and D147N mutant) were transfected again with the pKT3-Ub-R-GFP plasmid and dual transfectants were selected with 9µg/ml hygromycin B and G418. The concentration was gradually increased to 40µg/ml for hygromycin B and 50µg/ml G418. Same procedure was followed to generate ameba dually transfected with pKT3-Flag-CSN2 and pEhHYG-tetR-O-CSN5 overexpression plasmids.

### Cell viability assays

Parasite growth was measured using CellTiter-Glo® Luminescent Cell Viability Assay kit (Promega) (*11*). CSN5 overexpression and knockdown was induced by treating transfected parasites with 10µg/ml tetracycline, 0.1%DMSO. Empty vector control cells were treated the same conditions. Parasites of respective genotypes were plated in flat black bottom 96 well plates (Corning™ 96-Well Solid Black Polystyrene Microplates), 2500 cells/well in 300µl TYI-S-33 medium containing tetracycline. Plates were incubated at 37 °C in anaerobic chambers with GasPak™ EZ Gas Generating Container System (BD Diagnostics) to maintain the anaerobic condition during the incubation for indicated time points. At each time point plates were equilibrated to room temperature for 10 minutes followed by careful removal of media and replacing with 100 µl 1XPBS and 100µl CellTiter-Glo. Cell lysis was induced by placing the plates on an orbital shaker at RT for 10 min and then equilibrated at RT for 10 min to stabilize the luminescent signal. ATP-bioluminescence of the trophozoites were measured at RT using Synergy HTX Multi-Mode Reader (BioTek). For drug dose response assays 10,000 cells per well were cultured in medium containing either ZnDTC (.01nM −10µM, 0.1% DMSO) or metronidazole (.01nM-1mM, 0.1%DMSO), or 0.1% DMSO. Viability of parasites was measured by ATP-bioluminescence 48 hours after incubation. The effect of CSN5 overexpression and knockdown on drug susceptibility was measured and compared with the empty vector control cells. All three genotypes were induced with 10µg/ml tetracycline 6 hours before plating them for the assay (10,000 cells per well) in media containing 10µg/ml tetracycline and increasing doses of ZnDTC (0, 12.5, 25 and 50 nM, 0.1% DMSO).

### Protein degradation studies

Parasites transfected with the UPS substrate Ub-R-GFP protein degradation reporter (*44*) and either the mutant CSN5 expression plasmid or the pEhHYG-tetR-O-CAT empty vector control were treated with 10µg/ml tetracycline for 16 hours before confocal microscopy. Parasites harboring only the reporter plasmid were used for assaying protein degradation upon ZnDTC treatment. Cells were treated with 1µM ZnDTC or 0.1% DMSO control for 12 hrs prior to microscopy. Parasites were prepared for imaging by removing the media, pelleting by spinning at 200g for 5 minutes and resuspending cells in 1XPBS immediately before imaging. Confocal micrographs were analyzed and mean fluorescence intensities were calculated using Image-J software. For direct quantification of fluorescence, resuspended cells were distributed at a density of 20,000 cells/well in 100µl 1xPBS in black bottom 96 well plates (Corning™ 96-Well Solid Black or White Polystyrene Microplates). Fluorometric assay (*44*) was performed by measuring GFP fluorescence using Synergy HTX Multi-Mode Reader (BioTek). For the ZnDTC dose response experiment cells were treated in flasks with different doses of ZnDTC or 0.1%DMSO for 16 hours followed by fluorometric assay.

### Antibody purification

Purified GST-CSN5, GST-cullin1 and MBP-Nedd8 were used to immunize chicken, rabbit and guinea pigs (Cocalico Biologicals), respectively, and pre-immune serum, test bleeds, and the final bleed were received and tested by Western blotting. Sera were cleared of anti-GST or anti-MBP and non-specific antibodies prior to affinity purification with respective antigens. Affinity pull down of specific antibodies were done at 4°C overnight, using antigens conjugated to NHS-activated agarose beads (Pierce™ NHS-Activated Agarose Spin Columns, Thermo Fisher Scientific). Antibodies were eluted in 0.1M glycine, pH 2.8 and neutralized with 1M Tris, pH 9.0, and buffer exchanged into PBS with a 30kDa MWCO Centrifugal Filter (EMD Millipore).

### Immunoprecipitation and immunoblotting

Immunoprecipitations were performed with amebic lysates prepared from parasites of following genotypes: CSN5 knockdown, CSN5 overexpression, Flag-CSN2 overexpression, empty vector controls and untransfected wildtypes either treated with 1µM ZnDTC or 0.1% DMSO. Ameba were harvested and lysed with IP Lysis buffer (Thermo Fisher) supplemented with complete Protease inhibitors (Roche) and 1 mM PMSF (Roche). Lysate was sonicated and briefly cleared at 800g for 5 minutes. Total protein was measured via Bradford Assay (Bio-Rad). For CSN2 Co-IP, rabbit Anti-Flag magnetic agarose (Pierce) were incubated with the lysate overnight at 4°C. Agarose was washed, and bound protein was eluted with 1.5 mg/mL 3 X Flag peptide according to manufacturer’s directions. CSN5 Co-IP was performed with chicken anti-E. histolytica CSN5 antibody (5µg) using the Pierce™ MS-Compatible Magnetic IP Kit, streptavidin following manufacturer’s protocol. For cullin1 and Nedd8 pulldown, rabbit anti-*E. histolytica* cullin1 antibody (5µg) or guinea pig anti-*E. histolytica* Nedd8 antibody (5µg) conjugated to Dynabeads™ Protein A or Protein G, respectively, were used. 300µg total protein was used for each pulldown. For immunoblotting, samples were resolved on 4-20% polyacrylamide gels and transferred to nitrocellulose membrane (Bio-Rad). Membranes were blocked with 3% BSA or 5% milk and probed with the following primary antibodies-1:1000 rabbit anti-Flag (Cell Signaling technology); 1:100 chicken anti-*E*.*histolytica* CSN5 (custom); 1:250 rabbit anti-*E*.*histolytica* cullin1(custom); 1:500 guinea pig anti-*E*.*histolytica* Nedd8 (custom). Primary antibodies were detected by the following secondaries-1:10,000 goat anti-rabbit IgG HRP (Sigma Aldrich); 1:5000 donkey anti-chicken IgG HRP (Sigma Aldrich) and 1: 5000 Goat anti-guinea pig IgG HRP (Abcam).

### Mass spectrometry analysis

Immunoprecipitated proteins from the anti-Flag pulldown and anti-*E. histolytica* CSN5 were analyzed by mass spectrometry as described previously (*16*). The samples were processed by the W. M. Keck Biomedical Mass Spectrometry Laboratory at the University of Virginia. Label-free quantification was performed using MaxQuant (*45*).

### Parasite infection and treatment

Wild-type CBA/J mice at 8-10 weeks of age were obtained from the Jackson Laboratory. Mice were infected by intracecal injection of pHTP.luc plasmid transfected *E. histolytica* trophozoites (*40*), and vector expression was maintained with 50 µg/mL neomycin in drinking water. Treatment began 24 hours after infection (*11*), confirmed first by in vivo bioluminescent imaging and continued for total 5 days. The treated group were given 50 mg/kg disulfiram (Sigma Aldrich) and 1 mg/kg zinc gluconate (Alfa Aesar) suspended in Ora-plus (Paddock laboratories) via oral gavage. Untreated control groups were given Ora-plus only. Experiments were repeated with disulfiram and zinc gluconate from MedChemExpress and Pure Encapsulations respectively. Zinc gluconate was chosen over other formulations because of its safety profile. Dosing was based on previous animal studies that simulate the drug concentrations and pharmacokinetics of persons on FDA-recommended doses (*32, 35, 46*).

### *In vivo* bioluminescent imaging

Mice infected with luciferase-expressing parasites were injected IP with 150 µL Rediject Luciferin (Perkin Elmer) and imaged by the Xenogen IVIS II System.

### Histology and Immunohistochemistry

Mouse cecal tissue, fixed in Bouin’s solution (Sigma) and stored in 70% ethanol, was processed and stained with hematoxylin and eosin by the University of Virginia Research Histology Core. Histological scoring was performed by 2 independent blinded scorers and carried out as previously described (*40*). Mouse immunohistochemical staining was done using the DAKO Autostainer Universal Staining System with antibody against *E. histolytica* MIF protein (*10*).

### ELISA

Intestinal tissue lysates were evaluated by ELISA for myeloperoxidase (MPO) (R&D Systems) (*40*), total protein concentration was measured using the Pierce™ BCA Protein Assay Kit (Thermo Scientific).

### Bioinformatic analyses

Amino acids from protozoan CSN5 metalloprotease site and CSN2 PCI domains were aligned by Multiple Sequence Comparison by Log Expectation (MUSCLE) software. Structural modelling of *E. histolytica* CSN5 protein was done by Protein Homology/Analogy Recognition Engine v 2.0 (PHYRE^2^).The structures were visualized and analyzed using the UCSF Chimera software v. 1.10.2. Virtual screening was performed with drug molecules for their ability to dock with the CSN5 catalytic site using Autodock 4.2 (*47, 48*).

### Statistical analyses

Statistical differences were determined using Fisher’s exact test, Mann–Whitney *U* test and two tailed *t* test. Pearson’s correlation was used for correlation analysis. Nonlinear robust regression analysis was performed on the drug dose response curves. A *P* value less than .05 was considered statistically significant.

## Acknowledgments

We thank William Petri, Nicholas Sherman, Stuart Berr, Pat Pramoonjago, Girija Ramakrishnan and the University of Virginia Mass Spectrometry Research Facilities, Molecular Imaging Core, Research Histology, and the Biorepository and Tissue Research Facilities.

## Funding

This project was supported by National Institutes of Health (NIH) R01AI026649-S1, K08AI119181, UVA seed grant, and the Robert Wood Johnson Foundation– Harold Amos Medical Faculty Development Program Award. The funders had no role in study design, data collection and analysis, decision to publish, or preparation of the manuscript.

## Author contributions

S.G., and S.M designed the study. S.G., L.F., A.S., L.L., J.P., and S.M. performed experiments. S.G., L.F., A.S., L.L., J.P., D.S., and S.M. analyzed data. S.G., L.F., and S.M wrote the manuscript.

## Competing interests

The authors declare that they have no competing interests.

## Data and materials availability

All data associated with this study are available in the main text or the supplementary materials.

## SUPPLEMENTARY MATERIALS

**Fig. S1.**
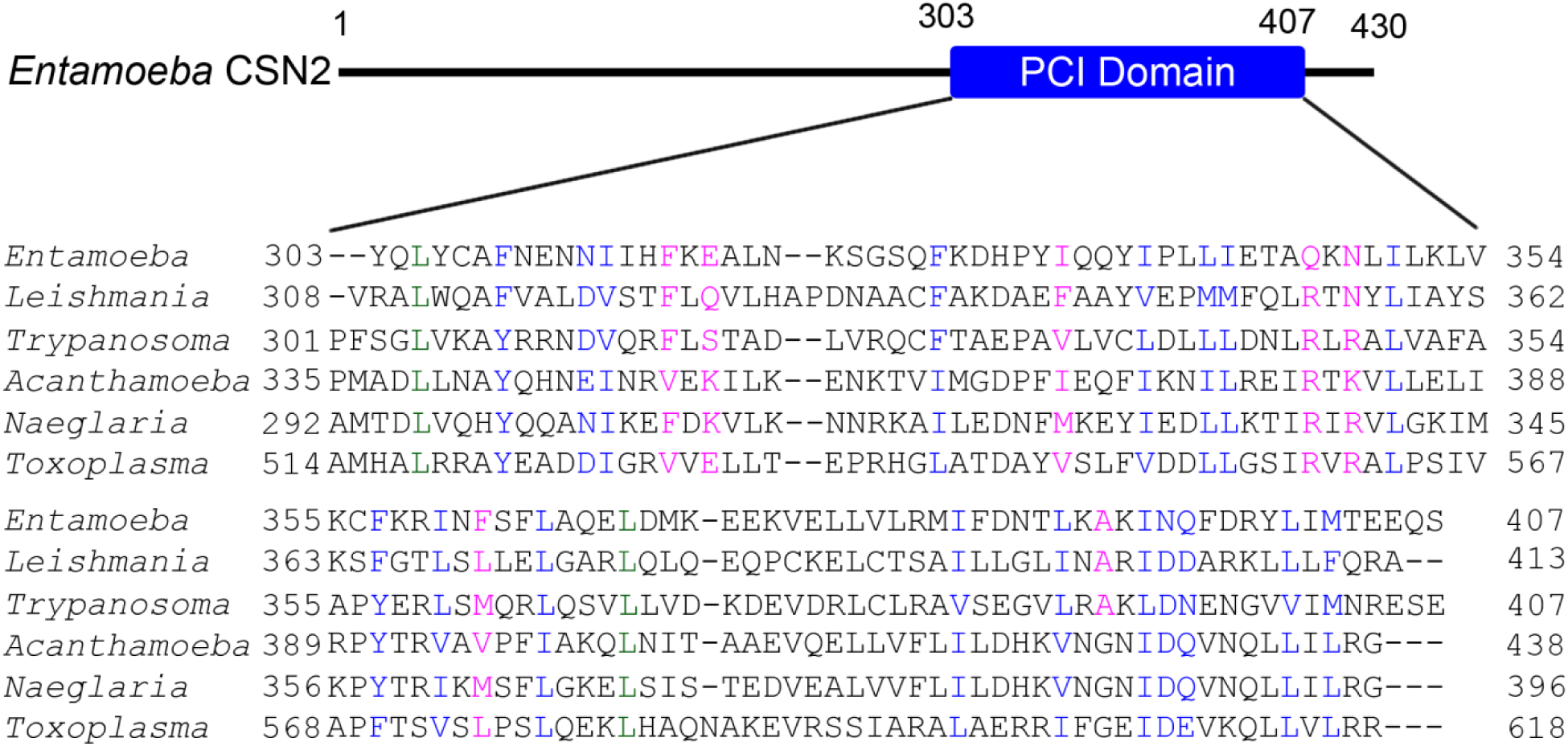
Alignment of the conserved PCI domain of CSN2 from pathogenic parasites. *Entamoeba histolytica* (EHI_174890A), *Leishmania donovani* (LdCL250028900-t42), *Trypanosoma cruzi* (Tb427.03.2320-t26), *Acanthamoeba castellanii* (ACAI_288500), *Naegleria fowleri* (Mrna1_nf0005120), *Toxoplasma gondii* (TGGT1_236220). Identical (green), conserved (blue), semi-conserved (pink), and non-conserved residues (black).

**Fig. S2.**
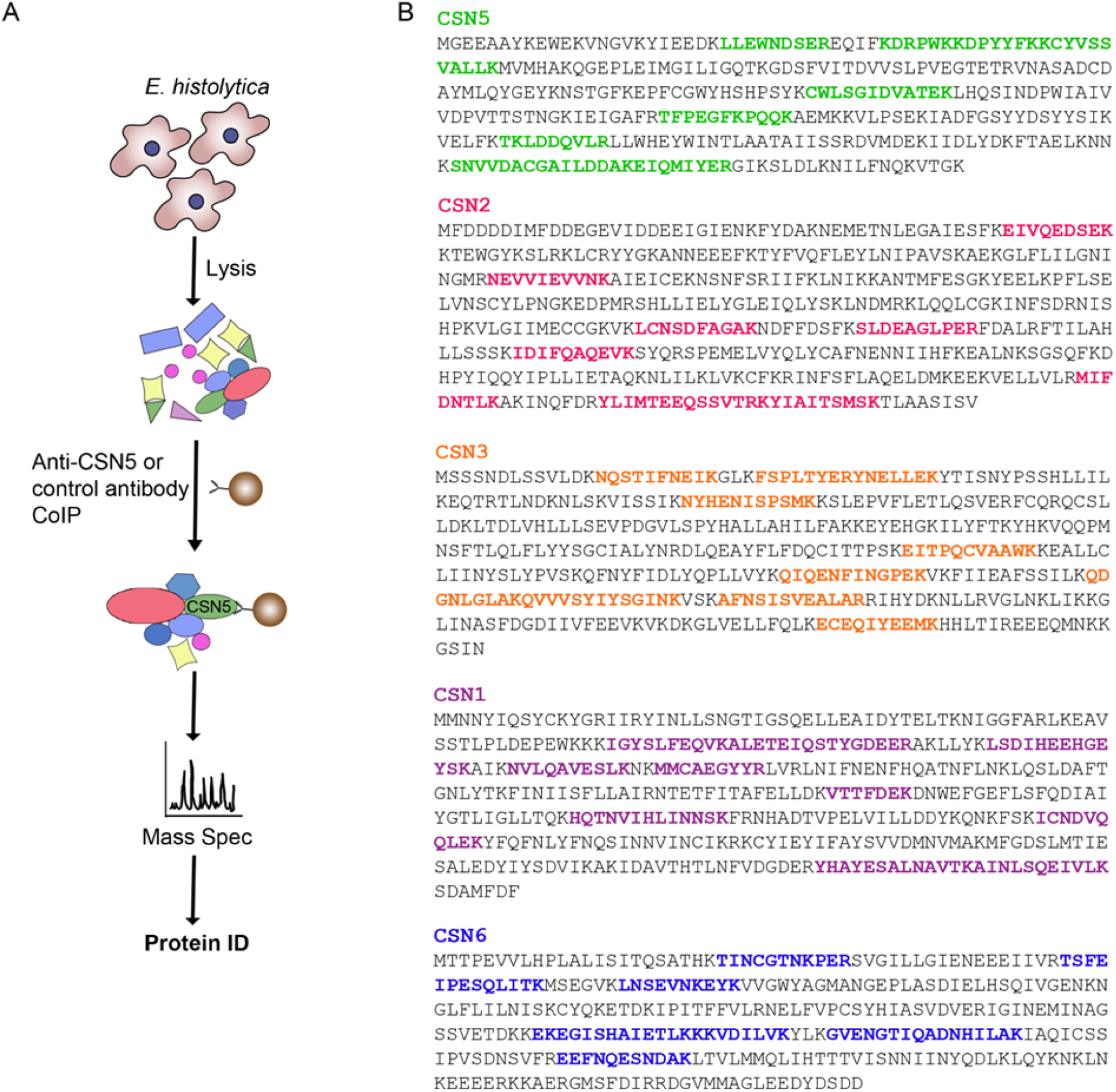
Parasite-encoded CSN subunits. (A) Schematic of procedure for identifying protein-protein interaction using co-immunoprecipitation with specific anti-CSN5 or control antibodies followed by mass spectrometric analysis (color highlighted). (B) CSN subunits amino acid sequences. Peptides unique to the CSN subunits identified as the top proteins detected only in the presence of anti-CSN5 and absent with control antibody co-immunoprecipitation.

**Fig. S3.**
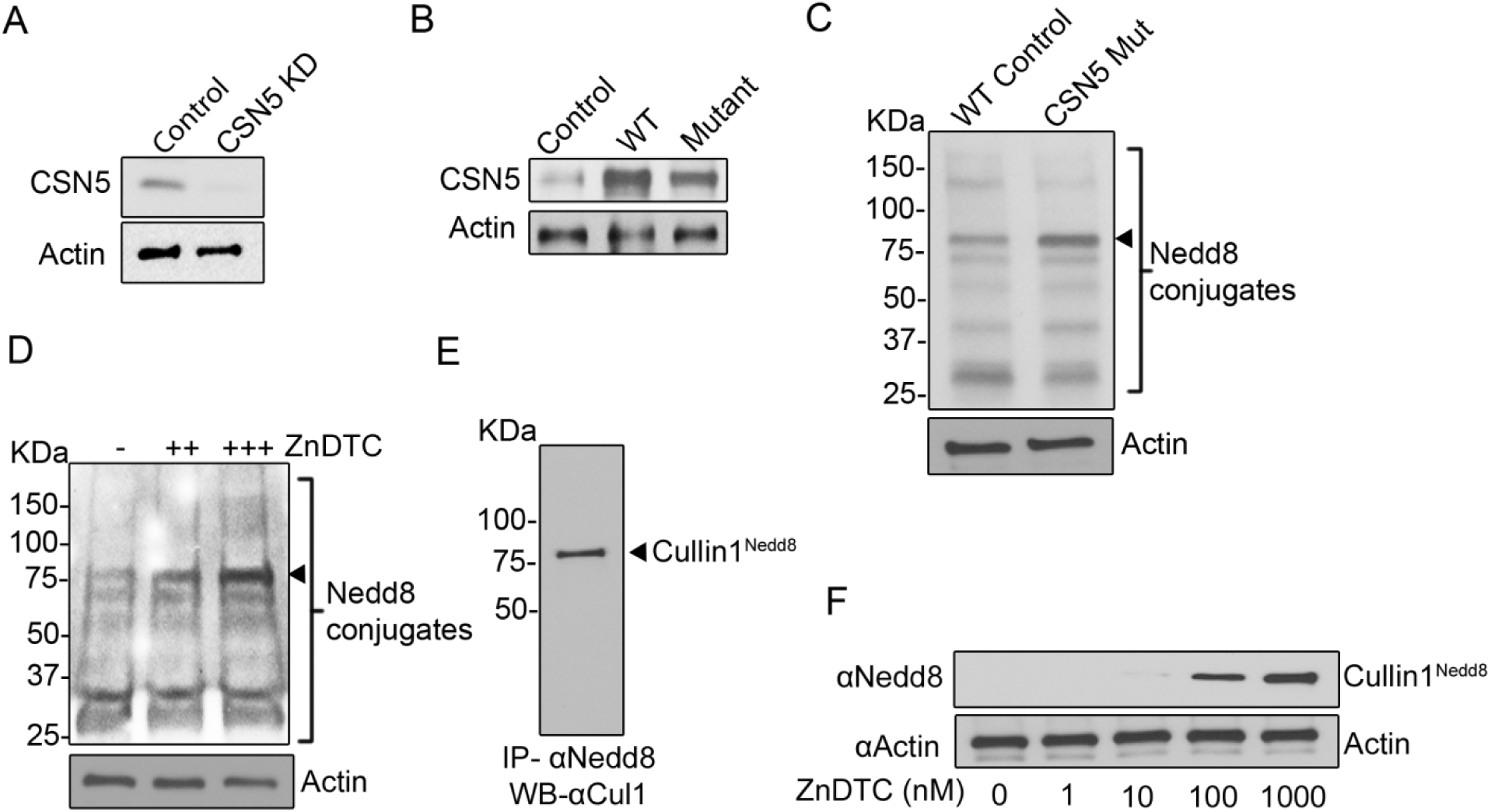
COP9 signalosome subunit 5 (CSN5) and neddylated protein expression in *E. histolytica* parasites. (A) Reduced CSN5 protein expression in knockdown cells compared to wild-type (WT) parasite controls with empty vector analyzed by immunoblotting (24 h). (B) Overexpression of WT and mutant (D147N) CSN5 proteins. (C and D) Accumulation of neddylated proteins in catalytically inactive CSN5 mutant and ZnDTC treated cells. (E) Immunoblot analysis neddylated cullin1. (F) Dose-dependent inhibitory effect of ZnDTC treatment on the endogenous deneddylation of cullin1. Actin used as loading control.

**Fig. S4.**
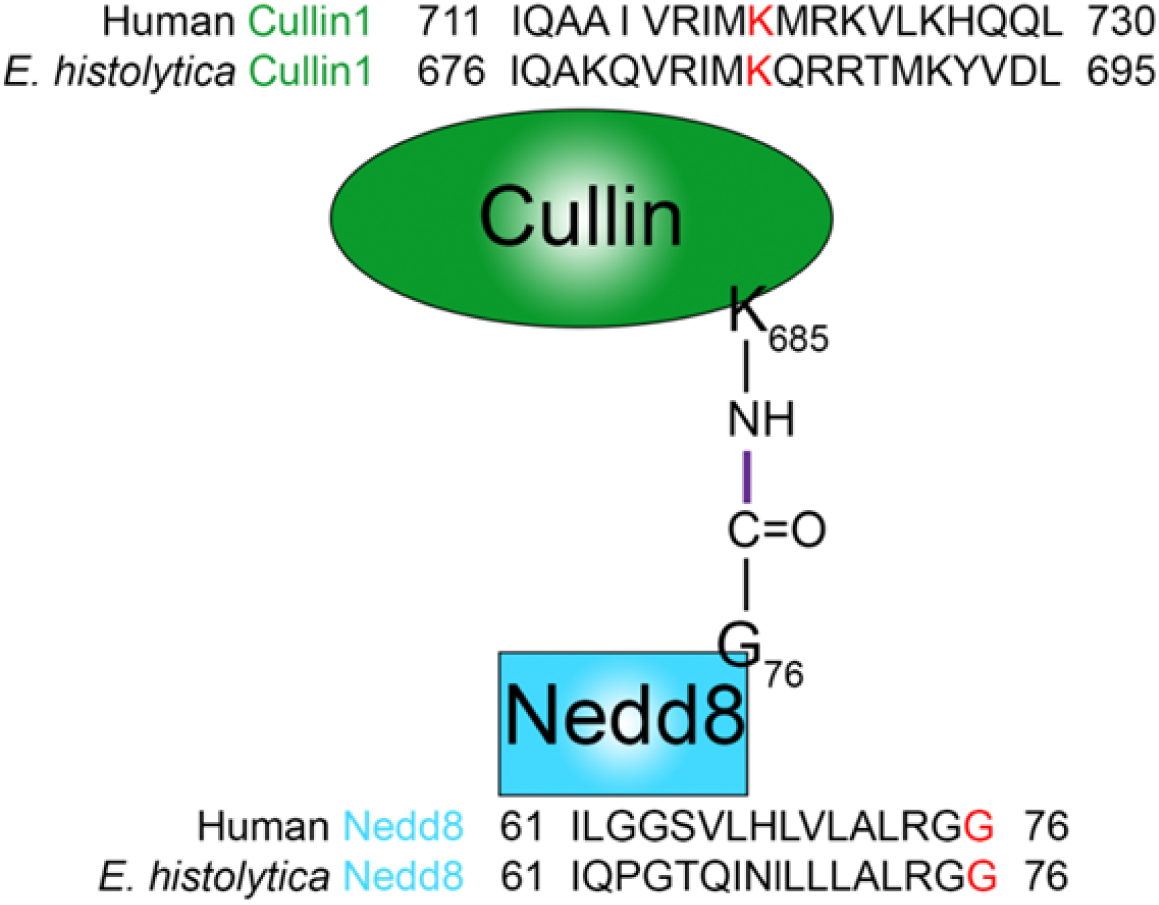
Neddylation, Nedd8 conjugated to cullin. *E. histolytica* cullin1 and Nedd8 showing isopeptide bond formation between the conserved lysine (K) residue of cullin1 and the C-terminal glycine (G) of Nedd8.

**Fig. S5.**
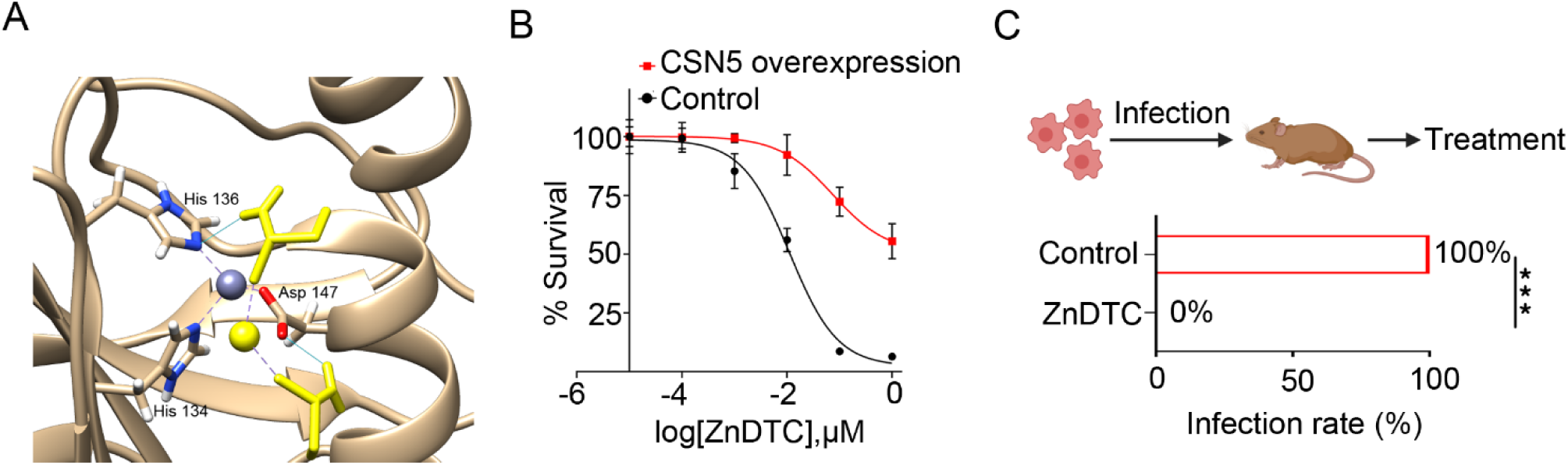
ZnDTC anti-parasitic activity. A) ZnDTC docks onto *E. histolytica* CSN5. Note the hydrogen bonds (blue lines) formed between ZnDTC drug (yellow) and the metalloprotease site Asp147 and His136. (B) Dose response curve showing increased resistance to ZnDTC treatment by *E. histolytica* parasites overexpressing CSN5. (C) Infection rate measured by ameba culture of cecal content, *n* = 7 mice per group. \*\*\**P* < .001, Fisher’s exact test.

